# Extracellular vesicles from non-neuroendocrine SCLC cells promote adhesion and survival of neuroendocrine SCLC cells

**DOI:** 10.1101/2022.10.12.511984

**Authors:** Lizandra Jimenez, Victor Stolzenbach, Patricia M. M. Ozawa, Marisol Ramirez-Solano, Qi Liu, Julien Sage, Alissa M. Weaver

**Affiliations:** Department of Cell and Developmental Biology, Vanderbilt University School of Medicine, Nashville, Tennessee.; The Center for EV Research, Vanderbilt University, Nashville, Tennessee.; Department of Biology, Northeastern University, Boston, Massachusetts.; Department of Biostatistics, Vanderbilt University Medical Center, Nashville, Tennessee.; Department of Pediatrics, Stanford University, Stanford, California.; Department of Genetics, Stanford University, Stanford, California.; Department of Pathology, Microbiology and Immunology, Vanderbilt University Medical Center, Nashville, Tennessee.

## Abstract

Small Cell Lung Cancer (SCLC) tumors are made up of distinct cell subpopulations, including neuroendocrine (NE) and non-NE cells. While secreted factors from non-NE SCLC cells have been shown to support the growth of the NE cells, the underlying molecular factors are not well understood. Here, we show that exosome-type small extracellular vesicles (SEVs) secreted from non-NE SCLC cells promote adhesion and survival of NE SCLC cells. Proteomic analysis of purified small EVs revealed that extracellular matrix (ECM) proteins and integrins are highly enriched in small EVs of non-NE cells whereas nucleic acid-binding proteins are enriched in small EVs purified from NE cells. Addition of select purified ECM proteins identified in purified EVs, specifically fibronectin, laminin 411, and laminin 511, were able to substitute for the role of non-NE-derived SEVs in promoting adhesion, survival, and tumorigenicity of NE SCLC cells. Those same proteins were differentially expressed by human SCLC subtypes. These data suggest that ECM-carrying SEVs secreted by non-NE cells play a key role in supporting SCLC tumor growth and survival.

## Introduction

Small Cell Lung Cancer (SCLC) is a highly aggressive cancer of neuroendocrine (NE) origin, characterized by a fast doubling time, early metastasis, and rapidly acquired therapeutic resistance ^1, 2^. Current treatments for SCLC make dual use of chemotherapy agents, such as etoposide and cisplatin in conjunction with targeted radiation ^3^. While immunotherapy has been shown to provide some effectiveness in a limited fraction of SCLC patients ^4^, treatment options remain limited and the 5-year survival rate is dismal at ~6% ^1^.

One of the factors that is thought to contribute to SCLC aggressiveness and treatment resistance is tumor heterogeneity. Both inter-tumor and intra-tumor heterogeneity is abundant in SCLC and is characterized by distinct NE and non-NE cellular phenotypes ^5, 6^. These distinct SCLC phenotypes have been classified in a variety of ways but are characterized by gene expression programs associated with key transcription regulatory factors, including the NE transcription factors ASCL1 and NEUROD1, and the non-NE transcription factors YAP1, and POU2F3 ^7–10^.

SCLC NE cells are thought to often originate from pulmonary NE stem cells, that sense oxygen levels and exist in a stem cell niche populated by non-NE cells ^11, 12^. This niche relationship may persist in SCLC tumors, that contain mixtures of cellular phenotypes including both NE and non-NE SCLC cells ^13^. Thus, in a transgenic mouse model of SCLC, a stem-like tumor-propagating cell population was identified that expresses high levels of NE markers and could self renew ^14^. In the same model, non-NE SCLC cells were shown to promote the growth of NE SCLC cells by non-cell autonomous secretion of factors, such as betacellulin and midkine ^15, 16^. Non-NE SCLC cells have also been shown to promote metastasis of NE SCLC cells ^17, 18^. This interaction between NE and non-NE SCLC cells is likely to be important for tumor cell survival in diverse environments, including in primary tumors, at metastatic sites, and in the context of therapy.

Secreted factors in the tumor microenvironment include both soluble factors and extracellular vesicles (EVs). EVs are membranous vesicles that carry a plethora of bioactive cargoes, including proteins, lipids, and nucleic acids. EVs are released from cells by diverse mechanisms, including secretion of exosomes from late endosomes and budding of ectosomes from the plasma membrane ^19^. EVs are involved in autocrine and paracrine communication and have emerged as key players in the tumor microenvironment ^19, 20^, interacting with diverse cell types to regulate growth, survival, angiogenesis, immune response, and metastasis. In SCLC, cancer derived EVs carrying the cytoskeletal protein profilin have been shown to promote cancer cell growth, angiogenesis, and lung metastasis ^21^. SCLC EVs can also have immunosuppressive effects, inhibiting CD8+ T-cells by carrying PD-L1 ^22^. In the brain, microvascular endothelial cell-derived EVs can promote SCLC survival, potentially contributing to the high rate of SCLC metastasis to the brain ^23^. Finally, EVs are being explored as a potential early diagnostic in SCLC ^24^.

In this study, we investigated the role of small EVs (SEVs) released from non-NE SCLC cells on the growth and phenotype of NE SCLC cells. We demonstrate that purified non-NE-derived SEVs promote growth, survival, and adhesion of NE cells. Quantitative comparative proteomics analysis of SEVs purified from non-NE and NE SCLC cells revealed enrichment of non-NE EVs with extracellular matrix proteins, suggesting candidate cargoes to promote growth and adhesion of NE cells. Tests of several of these cargoes identified fibronectin, laminin 411, and laminin 511 as potent regulators of NE SCLC growth and adhesion. These data indicate that specific ECM proteins carried by non-NE SCLC EVs support growth and adhesion of NE SCLC cells.

## Materials and Methods

### Antibodies

Rabbit anti-Tsg101 (cat no. ab30871), rabbit anti-Integrin alpha-2 (cat no. ab181548), rabbit anti-Collagen VI alpha 2 (cat no. ab180855), rabbit anti-Laminin gamma 1 (cat no. ab233389) mouse anti-Laminin alpha 4 (cat no. ab242359), rabbit anti-Laminin alpha 5 (cat no. ab184330), rabbit anti-Laminin beta 1 (cat no. ab184330), and rabbit anti-Tenascin C (cat no. ab108930) were purchased from Abcam. Mouse anti-Integrin alpha 5 (cat no. sc166665) were purchased from Santa Cruz. Rabbit anti-Integrin alpha 4 (cat no. 8440), rabbit anti-Integrin alpha 6 (cat no. 3750), rabbit integrin alpha v (cat no. 4711), rabbit anti-Thrombospondin-1 (cat no. 14778), rabbit anti-Gapdh (cat no. 5174), rabbit anti-Notch 2 (cat no. 5732), rabbit anti-Yap (cat no. 4912), rabbit anti-NeuroD1 (cat no. 7019), and rabbit anti-HES1 (cat no. 11988) were purchased from Cell Signaling. Rabbit anti-Fibronectin (cat no. F3648) was purchased from Sigma. Rabbit anti-Collagen 1 (cat no. ab765p) was purchased from Millipore. Rabbit anti-Fibulin-2 (cat no. GTX105108) was purchased from GeneTex. Mouse anti-Flotillin (cat no. 610820), mouse integrin beta 1 (cat no. 610467), and mouse anti-MASH1/ASCL1 (cat no. 556604) were purchased from BD Biosciences. Anti-Rabbit IgG (H+L), HRP Conjugate (cat no. W4011) and Anti-Mouse IgG (H+L), HRP Conjugate (cat no. W4021) were purchased from Promega.

### Tissue culture

The NE^KP3^ and non-NE^Hes1-GFP+^ mouse SCLC (mSCLC) cells were previously described ^15, 25^ and grown in RPMI-1640 medium supplemented with 10% bovine growth serum (BGS) (Cat no. SH30541.03, Hyclone) and penicillin–streptomycin (Cat no. 150-40-122, Gibco). For EVs collection, cells were culture using serum-free HITES medium ^26^ was prepared by adding 0.005mg/mL Insulin (Cat no.12585014, Gibco), 0.01 mg/mL Transferrin (Cat no. T5391, Sigma), 30 nM Sodium selenite (Cat no. S9133, Sigma), 10 nM Hydrocortisone (Cat no. H0135, Sigma), 10 nM Beta-estradiol (Cat no. E2257, Sigma), and 2 mM L-Glutamine Solution (Cat no. G7513, Sigma) to DMEM/F12 medium (Cat no. 10-092-CV, Corning). Human SCLC (hSCLC) cell lines NCI-H69 (A), NCI-H889 (A), NCI-H524 (N), NCI-H446 (N), NCI-H196 (Y) were grown in RPMI-1640 medium (cat no. 10-040-CV, Corning) supplemented with 10% fetal bovine serum (FBS) (cat no. F0926, Sigma). DMS53 (A2) cells were grown in Waymouth’s medium (cat no. 11220-035, Gibco) supplemented with 10% FBS. Finally, HITES medium prepped as stated above with the substitution with sodium bicarbonate DMEM/F12 (cat no. 11320-033, Gibco) and supplemented with 5% FBS was used to grow NCI-H841(Y) cells. All cells were cultured in incubators at 37°C and 5% CO2. For cell count we used Countess II automated cell counter (Thermo Fisher Scientific).

### Isolation of EVs from conditioned media

9 × 10^6^ NE^KP3^ and 2.4 × 10^6^ non-NE^Hes1-GFP+^ cells were plated in T182 culture flasks in RPMI growth media. The next day, the NE^KP3^ cells in the flasks were collected, washed three times with PBS, then resuspended in serum-free HITES media and added back to flasks. non-NE^Hes1-GFP+^ cells in flasks were washed three times with PBS before adding serum-free HITES media to the non-NE^Hes1-GFP+^ cells. After 48 h, the conditioned medium was collected from the cells and the EVs were isolated via serial centrifugation. Floating live cells and dead cell debris were removed from the conditioned medium after centrifugation steps of 300 × g for 10 min and 2,000 × g for 25 min, respectively. Large EVs were then collected by centrifugation at 10,000 × g for 30 minutes, followed by collection of the small EVs (SEVs) by centrifugation of the supernatant at 100,000 × g overnight. To quantitate the size and concentration of EVs, nanoparticle tracking analysis was performed using a Particle Metrix ZetaView PMX 110. The protein concentrations of total cell lysates were determined utilizing Pierce BCA Assay (Cat. 23225, Thermo Fisher Scientific). The protein concentrations of the EVs were determined utilizing Pierce Micro BCA Assay (Cat. 23235, Thermo Fisher Scientific).

### Conditioned media /EV transfer experiment

Non-NE^Hes1-GFP+^ conditioned media, non-NE^Hes1-GFP+^ conditioned medium depleted of EVs (aka Supernatant), and non-NE^Hes1-GFP+^ SEVs were isolated as described above. 1 × 10^6^ NE^KP3^ Cells were seeded in triplicate in a 6-well plate in 2 mL of media from one of 5 conditions: Conditioned Media (CM), Supernatant (Sup), serum-free HITES media, 75 × 10^6^ SEVs, 375 × 10^6^ SEVs. Cells were imaged via light microscopy (Nikon Eclipse TE2000E) at 48 and 72 hr post plating. After 72 hr, the total number of adherent and suspension cells in each well were counted and totaled as follows: **<u>For suspension cells:</u>** Media from each well was transferred to a 15 mL conical and the well was washed with 1 mL of PBS to collect any cells that may have been left behind. Each conical was then spun for 5 min at 100 × g. Supernatant was aspirated off and 500 μL of diluted (1:5) TrypLE Express (Cat no. 12604-013, Gibco) was added to each conical to dissociate cell pellet. After 2 min, we quenched TrypLE with 2.5 mL RPMI (1% P/S, 10% BGS). 10 μL of the quenched cells were taken for counting via trypan blue. **For adherent cells**: After the initial media was aspirated off for collection of suspension cells, we washed the well with 1 mL of PBS while being careful not to disturb the cells. We then aspirated off PBS and added 500 μL of diluted TrypLE to detach cells from well. After 2 min, we quenched TrypLE with 2.5 mL RPMI (1% P/S, 10% BGS). 10 μL of the suspension was taken for counting via trypan blue. All experiments were done in triplicates.

### Extracellular Matrix (ECM) Protein Assay

To assess the contribution of various ECM proteins on NE^KP3^ cells, 5 × 10^5^ NE^KP3^ cells were plated in duplicate wells of 12 well plates (Cat no. 3513, Corning) were plated in HITES media in the following conditions: 10 μg/mL Fibronectin (Cat no. 1918-FN-02M, R&D Systems) coated, 25 μg/mL Fibronectin coated, 2 μg/mL Fibulin-2 (Cat no. 9559-FB-050, R&D Systems) coated, 2 μg/mL Fibulin-2 added directly, 5 μg/mL Tenascin-C (Cat no. 3358-TC-050, R&D Systems) coated, 10 μg/mL Tenascin-C coated, 10 μg/mL Tenascin-C added directly, 5 μg/mL Laminin-411 (Cat no. LN411-0501, BioLamina) coated, and 5 μg/mL Laminin 511 (Cat no. LN511-0502, BioLamina) coated. Coating of wells was done overnight at 4°C the day prior to plating. Additionally, conditions plated for comparative analysis include: Conditioned media (positive control), serum-free HITES media (negative control), 75 SEV:cell, and 375 SEV:cell. Cells were imaged at 72 hr post plating and then assayed for viability via trypan blue exclusion. All experiments were done in triplicates.

### Serial Passaging of NE Cells in Conditioned Media

NE^KP3^ cells were thawed in normal media (RPMI with 10% BGS and 1% P/S). 72 hours after thawing, 3×10^6^ NE cells were plated into a T75 flask containing 10 mL of either CM, serum-free HITES, or Supernatant (Sup). **Passaging of suspension NE^KP3^ cells:** Media was collected into a 50 mL conical and flask was washed with 5 mL PBS to collect remaining cells. Cells were pelleted at 100 × g for 5 min. Supernatant was aspirated off and pellet was washed with 5 mL of PBS and spun down at 100 × g for 5 minutes. Supernatant was aspirated off and 2 mL of diluted (1:5) TrypLe was added to the pellet to dissociate cells. After 2 min cells were quenched with RMPI (w/ 10% BGS and 1% P/S), 10μL of the suspension was taken and cells were assessed for number and viability via trypan blue exclusion. 3×10^6^ cells were then moved to a 15 mL conical and washed with 10 mL PBS and pelleted for 5 min at 100 × g. Supernatant was aspirated off and cells were resuspended in 10 mL of appropriate media before being plated into a new T75 flask. **Passaging of Adherent NE^KP3^ cells:** Media was collected and any cells in suspension were counted as mentioned above. After collecting media adherent cells were washed with 5 mL of PBS. PBS was aspirated off and 2 mL of diluted (1:5) TrypLe was added to the plate to detach cells. After 2 min were quenched with RPMI (with 10% BGS and 1% P/S), 10 μL of the suspension was taken and cells were assessed for number and viability via trypan blue exclusion. 3×10^6^ cells were then moved to a 15 mL conical and washed with 10 mL of PBS and pelleted for 5 min at 100 × g. Supernatant was aspirated off and cells were resuspended in 10 mL of appropriate media before being plated into a new T75 flask. All cells were carried out to six passages.

### Mass spectrometry

#### Lysis of EVs for proteomics experiment

Non-NE^Hes1-GFP+^and NE^KP3^ SEVs were solubilized by adding to an equal volume of 2x Lysis Buffer [200mM triethylammonium bicarbonate buffer (TEAB) (cat. T7408, Sigma), 600mM sodium chloride (cat no. S23020, Research Products International), 2% Triton X-100 (cat no. A16046, Alfa Aesar), and 1% Sodium deoxycholate (cat no. 106504, Millipore)]. These lysates were sonicated with the Diagenode Bioruptor Standard Waterbath Sonicator (model cat no. UCD-200) in ice cold water for 15 min (30 sec on and off). Cleared lysates were collected after centrifugation at top speed (13,500 rpm) for 10 min in a Sorvall Legend Micro 21R Centrifuge (Thermo Fisher Scientific). The protein concentration of the cleared lysates was determined by Pierce Micro BCA Assay (Cat. 23235, Thermo Fisher Scientific).

Protein samples were precipitated, reduced, alkylated, and digested with trypsin as described in Jimenez *et al.*^27^. Quantitative proteomics analysis was performed using Thermo Fisher Scientific TMT Isobaric Mass Tagging reagents. According to the manufacturer’s instructions (Thermo Fisher Scientific), peptides were labeled with TMT reagents with each reconstituted protein sample being labeled with an individual vial of 0.8 mg TMTsixplex™ reagent. The resulting labeled peptides were then desalted by a modified Stage-tip method, reconstituted in 0.1% formic acid, and analyzed via MudPIT as described in Jimenez *et al.* ^27^. Using a Dionex Ultimate 3000 nanoLC and autosampler, MudPIT analysis was performed with a 13-step salt pulse gradient (0, 25, 50, 75, 100, 150, 200, 250, 300, 500, 750mM, 1M, and 2M ammonium acetate). Following each salt pulse, peptides were gradient-eluted from the reverse analytical column and introduced via nanoelectrospray into a Q Exactive Plus mass spectrometer (Thermo Fisher Scientific). Mobile phase solvents consisted of 0.1% formic acid, 99.9% water (solvent A) and 0.1% formic acid, 99.9% acetonitrile (solvent B). For the first 11 SCX fractions, the reverse phase gradient consisted of 2–50% B in 83 min, 50% B for 2 min, and a 10 min equilibration at 2% B. For the remaining fractions, peptides were eluted from the reverse phase analytical column using a gradient of 2-98% B in 83 min, followed by 98% B for 2 min, and a 10 min equilibration at 2% B. The Q Exactive Plus was operated in the data-dependent mode acquiring HCD MS/MS scans after each MS1 scan on the 15 most abundant ions. Normalized collision energy was set to 31, dynamic exclusion was set to 25 s, and peptide match and isotope exclusion were enabled.

For identification of peptides, HCD tandem mass spectra were searched in Proteome Discoverer 2.1 (Thermo Fisher Scientific) using SequestHT for database searching against a subset of the UniprotKB protein database containing *Mus musculus* protein sequences. Default templates in Proteome Discoverer were used for processing and consensus workflows. Search parameters included trypsin cleavage allowing for two missed cleavage sites, carbamidomethyl (C) and TMTsixplex (K, N-terminus) as static modifications, and a dynamic modification of oxidation (M). Percolator validation was performed with a target false discovery rate (FDR) setting of 0.01 for high confidence identifications. For quantitative analysis, an average reporter S/N threshold of 10 was selected, and no scaling or normalization modes were applied. Results were filtered to include those proteins for which a minimum of two unique peptides were identified. Log2 protein ratios calculated in Proteome Discoverer were then fit to a normal distribution using non-linear (least squares) regression, and the mean and standard deviation values derived from the Gaussian fit of the ratios were used to calculate *p* values. Statistically significant changes were determined using the Benjamini-Hochberg method with alpha level 0.05 (<5% FDR) ^28^.

##### Panther analysis

PANTHER (www.pantherdb.org) was used to identify protein classes of the proteins enriched in non-NE^Hes1-GFP+^ SEVs and NE^KP3^ SEVs.

##### Gene set enrichment analysis

Proteins were ranked by the log2 ratio between NE^KP3^ derived SEVs and non-NE^Hes1-GFP+^ derived SEVs. Gene Set Enrichment Analysis was performed on the pre-ranked proteins against the curated C2 and Hallmark gene sets from the Molecular Signatures Database (MSigDB v7.1).

##### FunRich analysis

Proteins that were two-fold upregulated in non-NE^Hes1-GFP+^ SEVs were uploaded into the software FunRich to identify any protein-protein interactions.

##### Western blot analysis

For Western blots, the mouse and human SCLC cells were lysed in RIPA buffer (50mM Tris-HCl, pH 7.6, 150mM NaCl, 1% NP-40, 1% Sodium deoxycholate, 1% SDS) with 1x protease inhibitor cocktail (cat no. 04693159001, Roche). 15 μg of mSCLC TCLs and SEVs were boiled in 2x SDS-PAGE sample buffer (4x SDS sample buffer: 0.2M Tris, pH 6.8, 8% SDS, 40% glycerol, 0.16M DTT and 0.29M bromophenol blue in water) for 5 min and loaded on 15-well 9% polyacrylamide gels. For the hSCLC cells, 20 μg of hSCLC TCLs were prepared for loading for Western blotting. The electrophoresis ran for 1 h 30 min at 110 volts at Room Temperature (RT). Proteins were transferred to nitrocellulose membranes for 1 h at 100 volts at 4°C. Membranes were blocked in 5 % non-fat dry milk diluted in Tris-buffered saline with 0.5 % Tween 20 (TBST) for 1 hour at RT. Primary antibodies were diluted in 5% non-fat dry milk - TBST (Thrombospondin-1, 1:5000; Integrin alpha-2, 1:5000; Integrin alpha 4, 1:5000; Integrin alpha-5, 1:10000; Integrin alpha 6, 1:5000, integrin alpha v, 1:5000, integrin beta 1, 1:5000, Collagen 1, 1:5000; Collagen VI alpha 2, 1:5000; Laminin alpha 4, 1:5000; Laminin alpha 5, 1:5000; Laminin beta 1, 1:5000; Laminin gamma 1, 1:5000; Tenascin C, 1:5000; Fibronectin, 1:10000; Fibulin-2, 1:5000; Tsg101, 1:5000; and Gapdh, 1:10000) and incubated overnight at 4°C. Membranes were washed 3 times for 10 min in TBST and subsequently incubated with species-specific HRP-conjugated secondary antibodies (1:10000; Promega) in 5% non-fat dry milk -TBST for 1 h at RT. All membranes were washed 3 times for 10 min in TBST and incubated with an enhanced chemiluminescence (ECL) reagent (Thermo Scientific) for 1 min before being imaged on Amersham 680 imager (GE). The densitometry measurements for the protein bands were done using the Analyze Gels feature of ImageJ (NIH) ^29^.

##### Statistical analysis

Experimental data were acquired from at least three independent experiments. Most data were compared using Welsh t-test and plotted as min-max in a box and whiskers plot in GraphPad Prism 8.

## Results

### Non-NE SEVs enhance NE SCLC cell growth and adhesion

To elucidate the role of EVs and other secreted factors on SCLC growth and survival, we cultured NE SCLC cells in the presence of conditioned media or purified SEVs from non-NE SCLC cells for 72 hours. For these studies, we used mSCLC cell lines derived from well-characterized genetically engineered mouse models of SCLC, which recapitulate many of the aspects of hSCLC, including heterogeneous cell composition, aggressive tumor behavior, and molecular driver and tumor suppressor characteristics. As a model of NE SCLC, we used KP3 cells (NE^KP3^), that are derived from an *Rb/p53* mutant mouse tumors ^25, 30, 31^. As a model for non-NE SCLC cells, we used Hes1-GFP+ non-NE (non-NE^Hes1-GFP+^) cells derived from a *Rb/p53/p130* mutant mouse in which GFP is expressed from the *Hes1* gene promoter, a target of Notch signaling and a marker of the non-NE state^15^.

To produce conditioned medium, non-NE^Hes1-GFP+^ cells were cultured in the presence of serum-free HITES medium ^26^ for 48 hr. To isolate SEVs, differential centrifugation of the conditioned media was performed with sequential pelleting of cells, debris, LEVs and SEVs at 300 × g, 2000 × g, 10,000 × g, and 100,000 × g and their number and size were characterized by nanoparticle tracking analysis **(Supplemental Figure 1 A).** Calculation of the EV numbers secreted by a known number of cells over 48 h revealed that non-NE^Hes1-GFP+^ cells secrete roughly 4 times more SEVs per cell compared to NE^KP3^ cells **(Supplemental Figure 1 B).** Very few large EVs were isolated; therefore, we only characterized and studied SEVs. When conditioned medium from non-NE^Hes1-GFP+^ cells was added to NE^KP3^ cells, the NE^KP3^ cells converted from growing in suspension to adhering to tissue culture plates **(Figure 1 A)**. This same adhesive morphological change was observed when unconditioned serum-free HITES media was supplemented with SEVs isolated from non-NE^Hes1-GFP+^ cells, in a concentration dependent manner **(Figure 1 A)**. Thus, while supplementation of medium with 50 SEVs per NE^KP3^ cell had no effect on NE^KP3^ morphology, ≥75 SEVs per NE^KP3^ cell induced adhesion. When the NE^KP3^ cells were grown in unconditioned medium or in conditioned medium depleted of EVs by ultracentrifugation (supernatant), the NE^KP3^ cells continued to grow in suspension **(Figure 1 A)**. Analysis of the cell number and viability revealed that the growth and survival of NE^KP3^ cells was also significantly enhanced by either treatment with conditioned medium or treatment with ≥75 non-NE SEVs per NE^KP3^ cell, compared to unconditioned serum-free HITES media or supernatant treatments **(Figure 1 B, C)**. The supernatant treatment also significantly reduced NE^KP3^ cell growth and viability compared to the unconditioned medium treatment **(Figure 1 B, C)**. Assessment of the adhesive phenotype by counting cells already in suspension versus those counted after trypsinization of adherent cells revealed an all-or-none shift with >99% of the cells cultured with conditioned medium or with ≥75 non-NE SEVs per NE^KP3^ cell adherent to the culture plate **(Figure 1 D)**.

**Figure 1:**
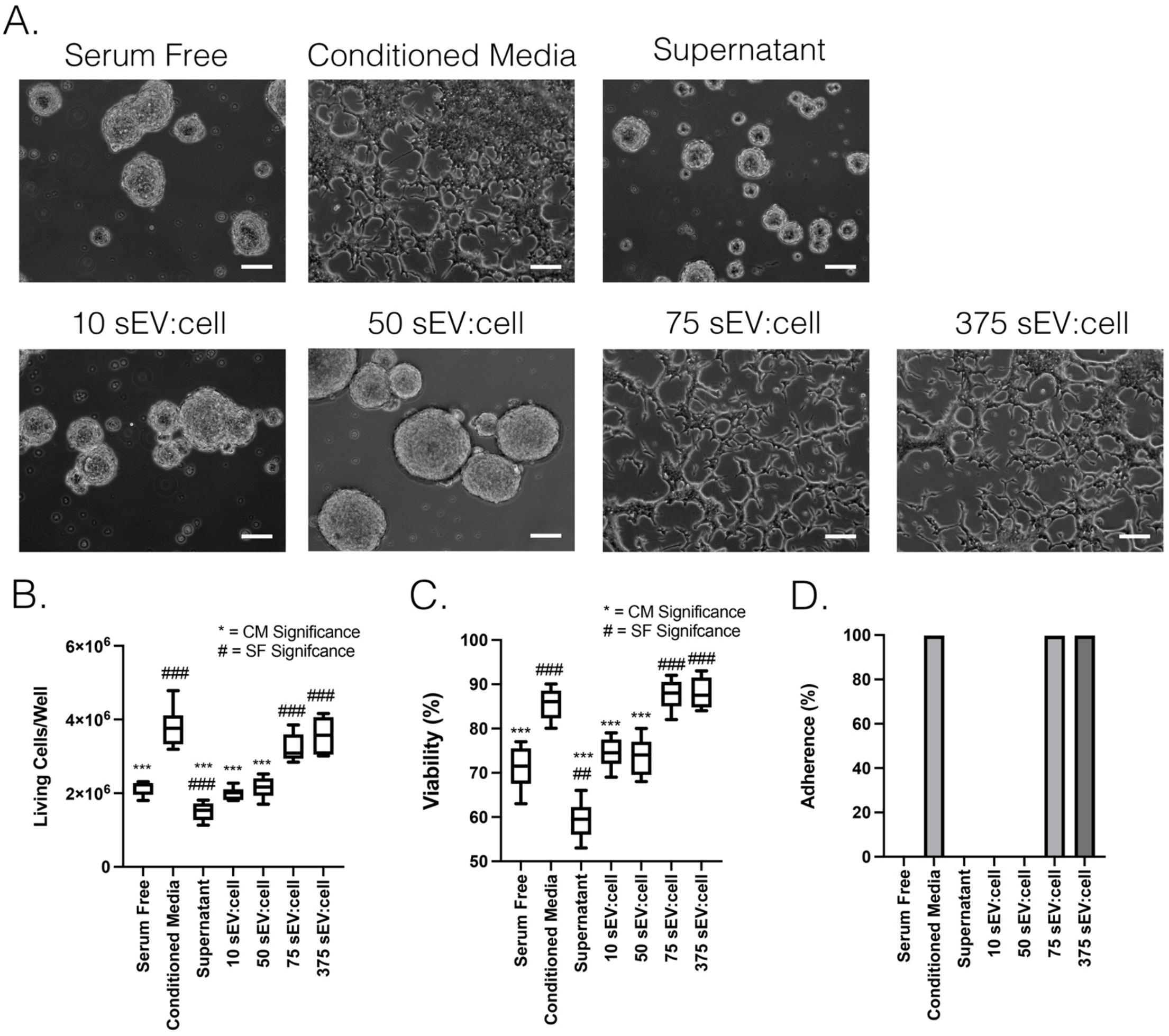
Non-NE^Hes1-GFP+^ SEVs enhance NE SCLC growth whilst eliciting an adhesive morphological change. A. Conditioned medium from non-NE^Hes1-GFP+^cells was added to NE^KP3^ cells before and after (Supernatant) ultracentrifuging overnight to remove EVs. Conditioned medium caused NE^KP3^ NE cells to shift to an adherent phenotype that was not seen when cells were cultured in Supernatant or unconditioned Serum Free HITES medium. Add SEVs purified from non-NE^Hes1-GFP+^ cells at four concentrations (10 × 10^6^, 50 × 10^6^, 75 × 10^6^, or 375 × 10^6^ SEVs) also induced an adherent morphology. B. NE^KP3^ cell numbers in all conditions measured via trypan blue exclusion assay show the media containing non-NE^Hes1-GFP+^ SEVs enhance NE^KP3^ cell growth. Box and whisker plot for three independent experiments. Box and whiskers plots with box indicating 25^th^-75^th^ percentile, whiskers showing min-max, and line indicating median. Note: *** p<0.001 show significance to conditioned media (CM), while ^###^ p<0.001 show significance to Serum-free HITES media. C. The percentage of viable NE^KP3^ were determined from trypan blue exclusion assay in B. Box and whisker plot for three independent experiments. Box and whiskers plots with box indicating 25^th^-75^th^ percentile, whiskers showing min-max, and line indicating median. Note: *** p<0.001 show significance to conditioned media (CM), while ^###^ p<0.001 show significance to Serum-free HITES media. D. The prominence of adhesive NE^KP3^ cells were assessed in each condition. Adhesive cells are considered those unable to be washed off by PBS.

### Conditioned media from non-NE^Hes1-GFP+^ cells does not alter levels of key transcription factors in NE^KP3^ cells

A distinguishing feature between different SCLC subtypes is their morphology in culture, with adhesion of non-NE cells to tissue culture plates and growth of NE cells in suspension ^15^. One possible explanation for the shift of NE^KP3^ cells to an adhesive morphology in the presence of EVs or conditioned media from non-NE cells could be induction of a phenotypic shift to a non-NE SCLC subtype. To test whether this is the case, we carried out a longer term experiment in which NE^KP3^ cells were cultured over multiple passages in conditioned media or EV-depleted supernatant collected from non-NE^Hes1-GFP+^ cells, or serum-free unconditioned media. As in our short term experiments, cells cultured in conditioned media adopted an adhesive morphology **(Figure 2 A**). Cell lysates from NE^KP3^ cells exposed to each of those conditions were collected after each passage and analyzed by Western blot for transcription factors that characterize and drive specific SCLC subtypes ^7, 32^ **(Figure 2 B**). As expected, parental NE^KP3^ cells were positive for the NE transcription factor ASCL1 and negative for the non-NE transcription factor YAP1 and for NOTCH2, which is active in non-NE SCLC cells, confirming these cells as “A” subtype SCLC cells ^7, 14^. By contrast, non-NE^Hes1-GFP+^ cells had the opposite profile (negative for ASCL1 and positive for YAP1 and NOTCH2), confirming them as a “Y-like” subtype of SCLC cells^15^. Neither cell line was positive for NEUROD1, which was present in the H524 “N” cells run on the same blot as a positive control. Treatment of NE^KP3^ cells with conditioned media for up to six passages did not alter this profile, with continued expression of ASCL1 and absence of expression of YAP1, NOTCH2, and NEUROD1. These data suggest that the induction of cell adhesion by conditioned media collected from non-NE^Hes1-GFP+^ cells is not due to a phenotypic shift to another SCLC subtype. Consistent with that conclusion, replacing the conditioned media with unconditioned media for 72 h led to a reversion of NE^KP3^ cells from an adhesive to suspension morphology **(Figure 2 C)**.

**Figure 2:**
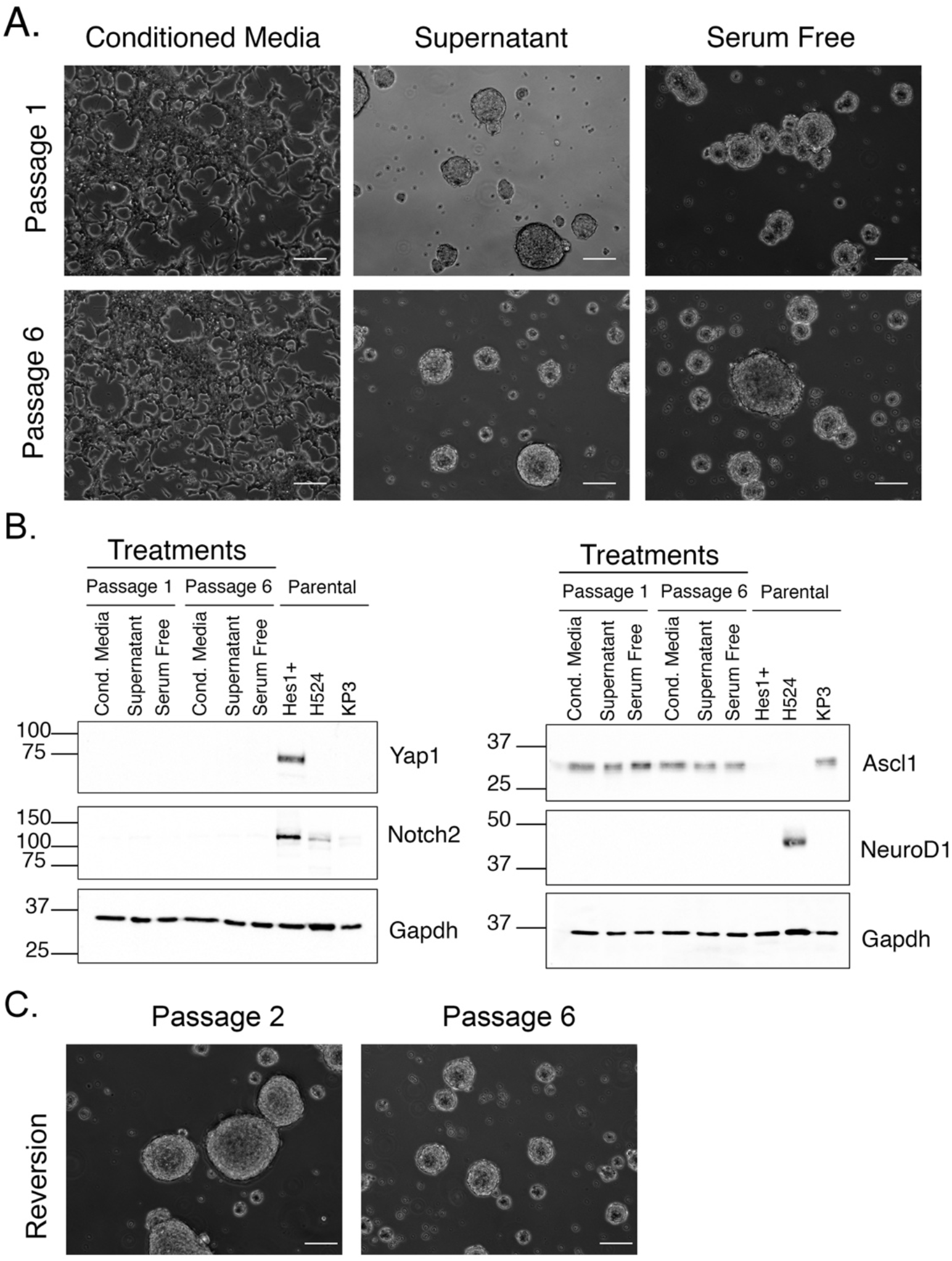
Serial passaging of NE cells in non-NE^Hes1-GFP+^ conditioned media enhances NE SCLC growth but does not elicit a permanent morphological change. A. Representative images of NE^KP3^ cells from passages1 and 6 in non-NE^Hes1-GFP+^ conditioned media, non-NE^Hes1-GFP+^ conditioned media devoid of extracellular vesicles (Supernatant), and unconditioned Serum-Free HITES medium. The adherent phenotype of NE^KP3^ cells grown in non-NE^Hes1-GFP+^ conditioned media is maintained for the six passages. Scale bar= 50 μM B. Western blot analysis of NE^KP3^ cells from passages 1 and 6 in the different conditions, as well as unconditioned parental cells, probing for marker proteins of different SCLC subtypes and GAPDH. As you can see, despite NE^KP3^ cells maintaining an adherent appearance, there was not an upregulation of marker proteins associated with the SCLC-Y (non-NE) and SCLC-N (NE NeuroD1) subtypes. C. Representative images that show that NE^KP3^ cells following passage 1 and 5 in non-NE^Hes1-GFP+^ conditioned media and replated in normal NE^KP3^ growth, have reverted to the suspension phenotype. Scale bar= 50 μM

### ECM proteins are an abundant protein cargo carried by non-NE small EVs

To assess what factors carried by SEVs purified from non-NE^Hes1-GFP+^ cells might promote adhesion and survival of NE^KP3^ cells, we isolated SEVs from each cell type and carried out relative and absolute quantitation of protein content by proteomics. Equal amounts of protein extracted from each EV type were labeled by tandem mass tagging and subjected to proteomics. The study reliably identified over 2300 proteins. Significantly upregulated proteins were determined by multiple comparisons using the Benjamini-Hochberg method. 194 and 27 proteins were significantly upregulated in the non-NE^Hes1-GFP+^ and NE^KP3^ SEVs, respectively **(Supplemental Table 1)**. In addition, the non-NE^Hes1-GFP+^ SEVs had 412 proteins identified to be upregulated by at least 2-fold (i.e. normalized fold change of 0.5 or lower) compared to the NE^KP3^ SEVs whereas NE^KP3^ SEVs had 138 proteins upregulated by at least 2-fold (i.e. ratio of 2 or higher) compared to non-NE^Hes1-GFP+^ SEVs **(Supplemental table 1)**.

**Table 1.** Abbreviated list of top GSEA gene sets. Some of the top gene sets for upregulated proteins in non-NE and NE SEVs. The non-NE^Hes1-GFP+^ SEVs were enriched with gene sets related to ECM organization and integrin-cell surface interactions. The NE^KP3^ SEVs were enriched with gene sets related to translation, RNA processing, and amino acid metabolism. For each gene set, the gene ontology (GO) name, # of proteins, normalized enrichment score, and false discovery rate (FDR) q value are listed.

|  | Geneset Name | # of proteins | Normalized Enrichment Score | FDR q value |
| --- | --- | --- | --- | --- |
| non-NE<sup>Hes1-GFP+</sup> SEVs | NABA Matrisome | 150 | -9.170 | 0 |
|  | REACTOME Extracellular Matrix Organization | 98 | -5.472 | 0 |
|  | REACTOME ECM Proteoglycans | 34 | -5.005 | 0 |
|  | REACTOME Degradation of The Extracellular Matrix | 42 | -4.957 | 0 |
|  | REACTOME Collagen Biosynthesis and Modifying Enzymes | 24 | -4.520 | 0 |
|  | PID Integrin1 Pathway | 34 | -4.442 | 0 |
|  | REACTOME Collagen Formation | 32 | -4.427 | 0 |
|  | REACTOME Assembly of Collagen Fibrils and Other Multimeric Structures | 23 | -4.212 | 0 |
|  | ZHONG Secretome of Lung Cancer and Endothelium | 49 | -4.210 | 0 |
|  | REACTOME Non-Integrin Membrane ECM Interactions | 30 | -4.118 | 0 |
|  | REACTOME Integrin Cell Surface Interactions | 31 | -4.072 | 0 |
|  | KEGG ECM Receptor Interaction | 34 | -4.052 | 0 |
|  | ZHONG Secretome of Lung Cancer and Fibroblast | 97 | -3.932 | 0 |
|  | PID Integrin3 Pathway | 19 | -3.886 | 0 |
|  | NABA ECM Glycoproteins | 55 | -3.867 | 0 |
|  | NABA Secreted Factors | 19 | -3.865 | 0 |
|  | REACTOME Collagen Degradation | 17 | -3.779 | 0 |
|  | ZHONG Secretome of Lung Cancer and Macrophage | 56 | -3.727 | 0 |
|  | NABA ECM Regulators | 33 | -3.698 | 0 |
|  | REACTOME Syndecan Interactions | 16 | -3.666 | 0 |
| NE<sup>KP3</sup> SEVs | KEGG Ribosome | 62 | 3.412 | 0 |
|  | REACTOME Response of EIF2AK4 GCN2 to Amino Acid Deficiency | 67 | 3.404 | 0 |
|  | REACTOME SRP Dependent Cotranslational Protein Targeting to Membrane | 65 | 3.378 | 0 |
|  | REACTOME Eukaryotic Translation Elongation | 68 | 3.371 | 0 |
|  | REACTOME Selenoamino Acid Metabolism | 72 | 3.308 | 0 |
|  | REACTOME Nonsense Mediated Decay | 78 | 3.177 | 0 |
|  | REACTOME Regulation of Expression Of Slits And Robos | 114 | 2.954 | 0 |
|  | REACTOME Eukaryotic Translation Initiation | 90 | 2.911 | 0 |
|  | REACTOME rRNA Processing | 101 | 2.893 | 0 |
|  | BILANGES Serum and Rapamycin Sensitive Genes | 50 | 2.836 | 0 |
|  | REACTOME Signaling by Robo Receptors | 132 | 2.759 | 0 |
|  | REACTOME Translation | 116 | 2.645 | 0 |
|  | REACTOME Metabolism of Amino Acids And Derivatives | 153 | 2.608 | 0 |
|  | BILANGES Serum Response Translation | 19 | 2.559 | 0 |
|  | REACTOME Metabolism of RNA | 323 | 2.067 | 0.002 |

Analysis of the data by protein class using Panther software revealed that a large fraction of the total proteins identified in non-NE^Hes1-GFP+^ SEVs were extracellular matrix (ECM) proteins (12.6%) and cytoskeletal proteins (8.4%), along with metabolite interconversion and protein modifying enzymes (23 and 12.3%) **(Figure 3 A)**. By contrast, for the NE^KP3^ SEVs the top two categories were translational protein (43%) and nucleic acid binding protein (18.3%) **(Figure 3 A)**. Conversion of the protein names to gene names and analysis of the data by gene set enrichment analysis (GSEA) further revealed multiple gene sets related to ECM and adhesion that are enriched in non-NE^Hes1-GFP+^ SEVs, including matrisome, ECM organization, ECM regulators, collagen formation, epithelial-mesenchymal transition, and integrin-cell surface interactions (selected gene sets shown in **Table 1** and **Supplemental Figure 2A**, full list of gene sets shown in **Supplemental Table 2**). By contrast, gene sets related to translation, ribosomes, RNA processing, and amino acid metabolism are enriched in NE^KP3^ SEVs. **(Table 1, Supplemental Figure 2B, and Supplemental Table 2)**. Finally, the software Funrich was used to identify protein-protein interactions in proteins upregulated in non-NE^Hes1-GFP+^ SEVs. This analysis identified a single main cluster centered around the ECM protein fibronectin, which is a key organizer of stromal ECM assembly **(Figure 3 B)**.

**Figure 3:**
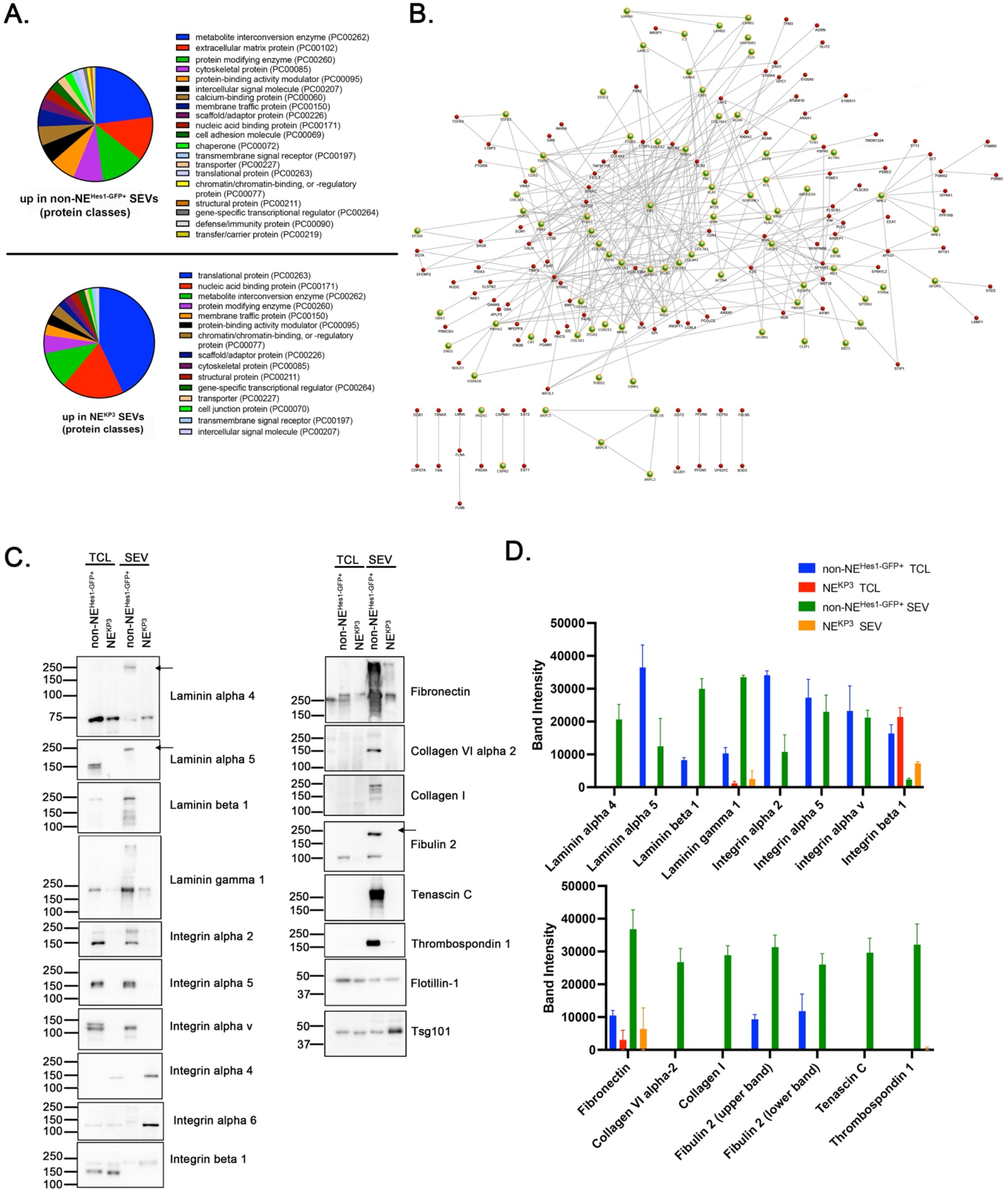
Identification of SEV protein cargo content that regulate different SCLC phenotypes. A. Panther protein classes of proteins showing elevated levels in non-NE^Hes1-GFP+^ (top) and NE^KP3^ (bottom) SEVs. B. FunRich Protein Interaction Network of the proteins that were two-fold higher in non-NE^Hes1-GFP+^ SEVs, highlighting the integrin family cell surface interactions with green dots. Of note, the main cluster seems centered around Fibronectin. C. Western blot analysis of non-NE^Hes1-GFP+^ and NE^KP3^ TCLs and SEVs probing for Laminins alpha 4 and 5, Laminin beta 1, Laminin gamma 1, Integrins alpha 2, 4, 5, 6, and v, and beta 1, Thrombospondin 1, Fibronectin, Collagen VI alpha 2, Collagen I, Tenascin C, Fibulin 2, and Tsg101. Arrows were added to Laminins alpha 4 and 5 and Fibulin 2 to indicate that these proteins running higher in non-NE^Hes1-GFP+^ SEVs. D. Quantitation of Western blots band intensities of candidate proteins.

To validate the proteomic data, Western blot analysis for selected proteins was performed on total cell lysates (TCLs) and SEVs from non-NE^Hes1-GFP+^ and NE^KP3^ cells **(Figure 3 C, D**). This analysis confirmed that the ECM proteins laminins alpha 4, alpha 5, beta 1, and gamma 1, fibronectin, and fibulin, along with integrins alpha 2, alpha 5, and alpha v were enriched in non-NE^Hes1-GFP+^ TCLs in comparison to NE^KP3^ TCLs. These same proteins were enriched in non-NE^Hes1-GFP+^ SEVs in comparison to NE^KP3^ SEVs **(Figure 3 C, D**, note that for some of the ECM proteins there is a shift to a higher mobility, possibly due to covalent crosslinking in assembled ECM). In addition, thrombospondin 1, collagen VI alpha 2, collagen I and tenascin C were exclusively found in the non-NE^Hes1-GFP+^ SEVs. Ponceau stains of the blots prior to blocking and antibody probing shows that the TCLs and SEVs were evenly loaded **(Supplemental Figure 3)**.

#### Select ECM proteins identified on non-NE SEVs promote NE cell growth and adhesion

The strong enrichment of ECM and adhesion proteins in non-NE^Hes1-GFP+^ SEVs suggests a likely mechanism for the functional activity of those SEVs in promoting adhesion and growth of NE^KP3^ cells. To test that possibility, specific purified ECM proteins identified to be enriched in non-NE^Hes1-GFP+^ SEVs were coated onto tissue culture plates and/or added into the media, followed by adding NE^KP3^ cells and culturing for 72 h. Thus, fibronectin, fibulin-2, tenascin-C, laminin-411, and laminin-511 were each tested. Fibronectin was chosen for testing due to its enrichment and identification as a protein-interaction hub in the FunRich analysis. Fibulin-2 is a regulator of TGFβ activity, that we had previously identified to be carried by astrocyte SEVs and promote neuronal synapse formation ^33^. Tenascin C is known as a metastatic niche protein in numerous cancers ^34–36^. Laminins 411 and 511 are known to promote normal stem cell growth and adhesion ^37–42^. Of these treatments, fibronectin, laminin 411, and laminin 511 all promoted NE^KP3^ cell growth and adhesion **(Figure 4 A-D)**. No other conditions exhibited an adhesive phenotype. Since Fibulin-2 and Tenascin C did not promote adhesion upon coating tissue culture plates, in some cases we added them into the media; however, they still did not promote adhesion of NE^KP3^ cells **(Figure 4 A, D)**. Only one Tenascin-C concentration showed any effect on cell growth, with a minor increase in cell numbers and viability. Interestingly, all Fibulin-2 conditions decreased cell numbers and viability **(Figure 4 B)**. Overall, purified Fibronectin, Laminin-411, and Laminin-511 all produced similar effects to purified EVs, suggesting that they may contribute to the EV-induced promotion of growth and adhesion.

**Figure 4:**
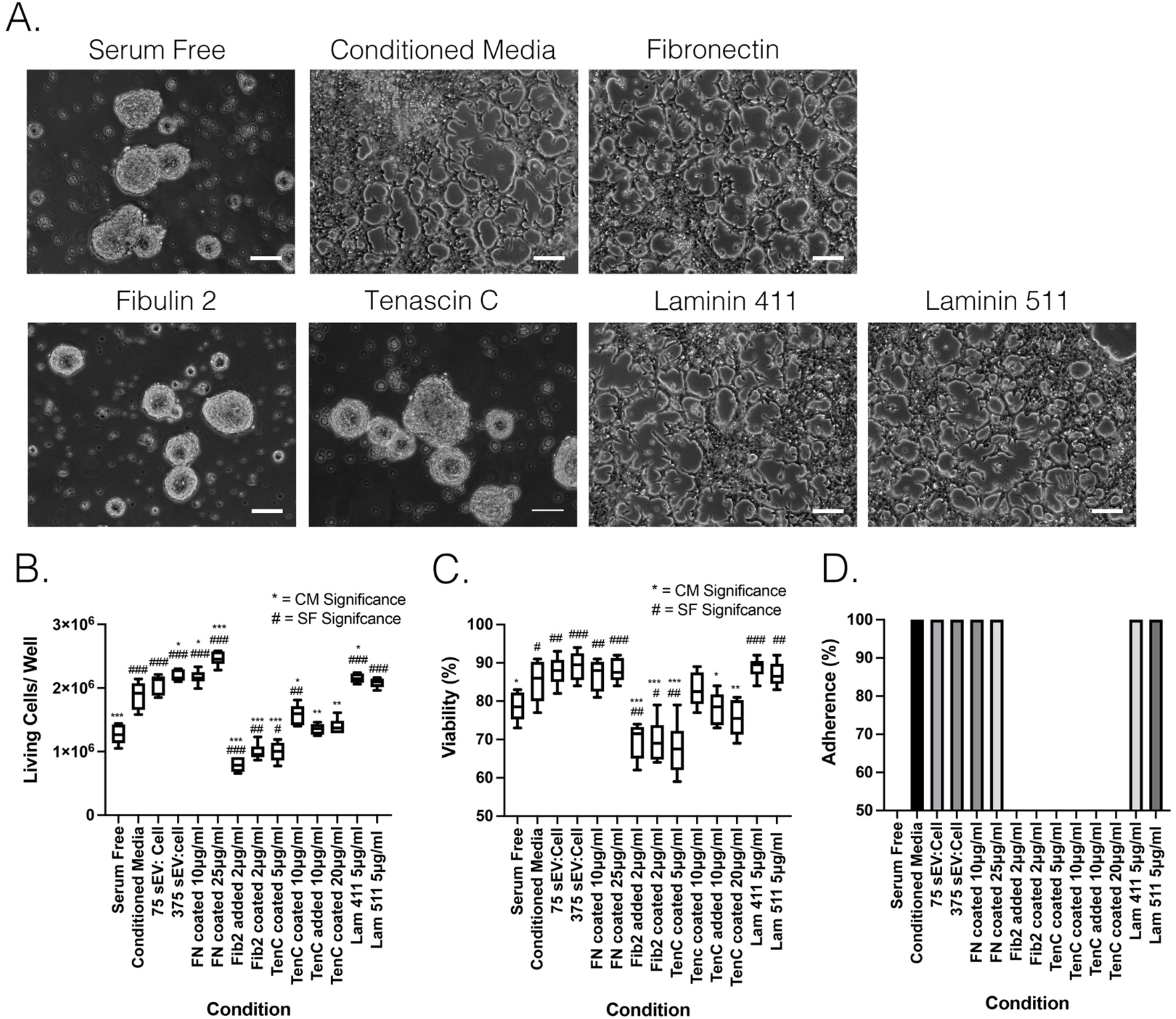
Select ECM molecules found in non-NE^Hes1-GFP+^ SEVs enhance NE SCLC growth and elicit a morphological change. A. Representative images of NE^KP3^ cells following ECM treatments. Coating with Fibronectin, Laminin 411 and 511 promoted an adherent phenotype, while fibulin 2 and tenascin-C did not. B. NE^KP3^ cell numbers in all conditions measured via trypan blue exclusion assay show that select ECM coating enhance NE^KP3^ cell growth. Box and whisker plot for three independent experiments. Box and whiskers plots with box indicating 25^th^-75^th^ percentile, whiskers showing min-max, and line indicating median. Note: * p<0.05, ** p<0.01, and *** p<0.001 show significance to conditioned media (CM), while ^#^ p<0.05, ^##^ p<0.01, and ^###^ p<0.001 show significance to Serum-free HITES media. C. The viability numbers for NE^KP3^ cells in all the conditions determined by the trypan blue exclusion assay. Box and whisker plot for three independent experiments. Box and whiskers plots with box indicating 25^th^-75^th^ percentile, whiskers showing min-max, and line indicating median. Note: * p<0.05, ** p<0.01, and *** p<0.001 show significance to conditioned media (CM), while ^#^ p<0.05, ^##^ p<0.01, and ^###^ p<0.001 show significance to Serum-free HITES media. D. Percent adherence of NE^KP3^ cells from all the conditions.

### Integrins and ECM proteins are differentially expressed by human SCLC subtypes

At least 4 different major SCLC subtypes have been identified, including A, A2, N and Y subtypes ^7, 32^. SCLC-A and SCLC-A2 cell lines express the NE driver transcription factor ASCL1 with SCLC-A cell lines growing as suspension cells and SCLC-Y cells express the mesenchymal transcription factor YAP1 and grow adherent to culture plates. SCLC-N cells express NEUROD1 and have both NE and mesenchymal/non-NE features ^43–46^ and can go as suspension (H524) of adherent (H446) cells. To determine whether the adhesion proteins we identified to be upregulated in the mouse non-NE “Y-like” subtype cells were also enriched in similar human SCLC cell lines, we carried out Western blot analysis. We first assessed transcription factors that are characteristic of different SCLC subtypes in various human SCLC cell lines. As expected, we detected ASCL1 in the two SCLC-A lines (H69 and H889), as well as the SCLC-A2 line (DMS53) (**Figure 5 A, B**). Both SCLC-N cell lines (H524 and H446) expressed NEUROD1. “Y” cell lines are characterized not only by expression of YAP1 but also by NOTCH activation, characterized by expression of NOTCH2 and the Notch-responsive gene HES1. Indeed, the two SCLC-Y cell lines (H196 and H841) express high levels of YAP1, HES1, and NOTCH2 **(Figure 5)**. NOTCH2 and HES1 expression was also identified in several other cell types. Notably, HES1 together with ASCL1 is expected in SCLC-A2 subtypes ^32^, which validates DMS53 as an SCLC-A2 cell type and suggests that H69 may be more of an SCLC-A2 subtype than a pure SCLC-A subtype **(Figure 5 A, B)**. NOTCH2 expression was also expressed in multiple cell types (**Figure 5 A, B)**. We also included our mSCLC cell lines on the blot and indeed NE^KP3^ cells express only ASCL1, suggesting that they are of the “A” subtype and non-NE^Hes1-GFP+^ cells express YAP1, HES1, and NOTCH2, as expected for a “Y-like” subtype.

**Figure 5.**
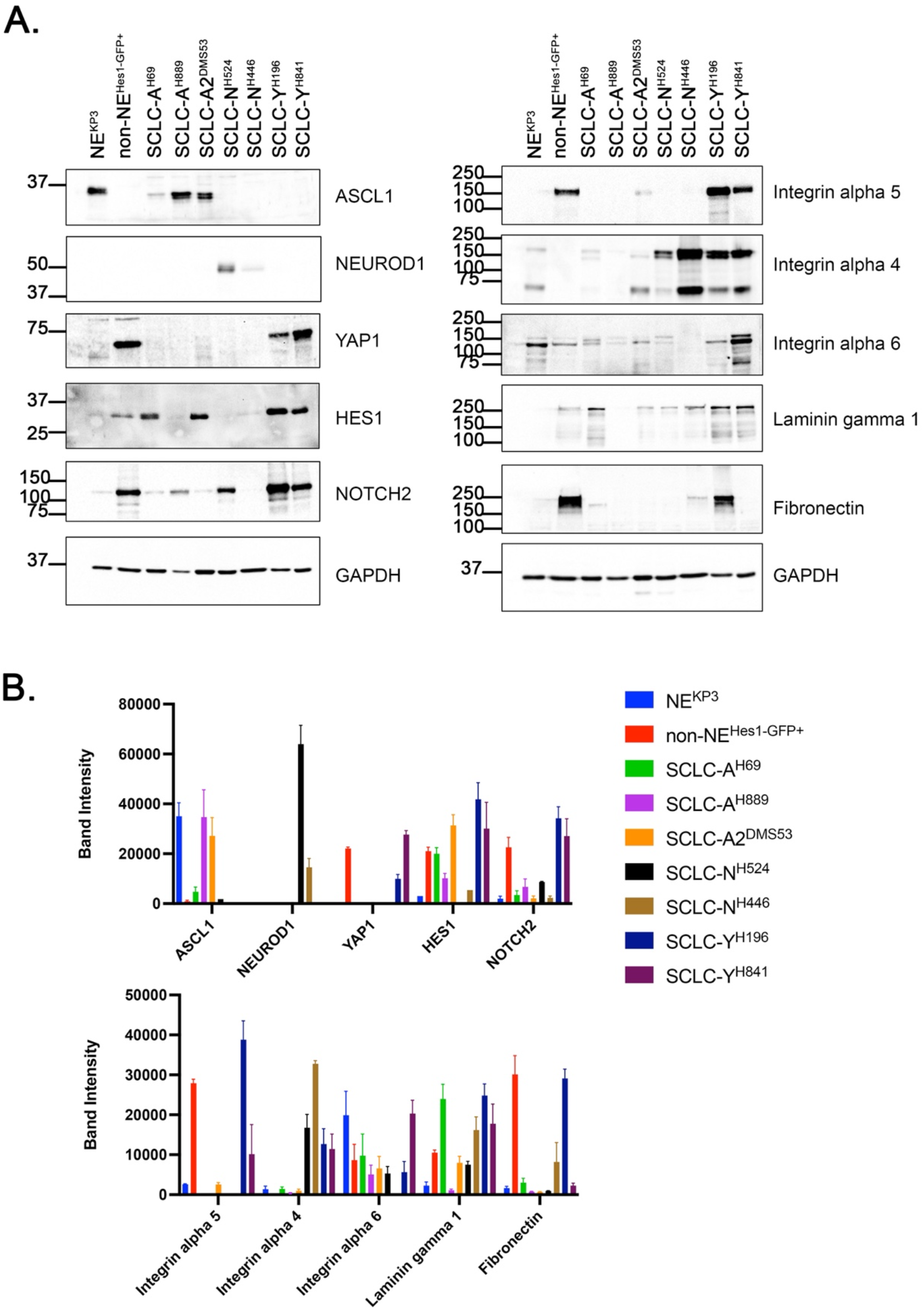
Analysis of integrin and ECM proteins expressed in human SCLC subtype cell lines. A. Western blot analysis of total cell lysates of human SCLC lines for transcription factors characteristic of SCLC subtypes (ASCL1, NEUROD1, and YAP1), integrins alpha 4, 5, and 6, and two ECM proteins (Laminin gamma 1 and Fibronectin). GAPDH was probed as loading control. B. Quantitation of Western blot band intensities of analyzed proteins.

We next examined expression in human SCLC cells of the integrins and ECM proteins identified to be differentially expressed by NE and non-NE mSCLC cells and EVs by Western blot analysis of cell lysates from both mouse and human SCLC cells. Similar to the mouse SCLC cells, integrin alpha 5 was greatly enriched in the two SCLC-Y cell lines and absent from the 2 SCLC-A cell lines. There was also a small amount of integrin alpha 5 expression in the SCLC-A2^DMS53^ cell line. In contrast to the mSCLC cell lines, integrin alpha 4 was more abundant in the SCLC-N and SCLC-Y lines than the SCLC-A lines and SCLC-A2 line **(Figure 5 A, B)**. As both integrin alpha 5 beta1 and integrin alpha 4 beta1 are fibronectin receptors, these data are consistent with more abundant fibronectin assembly by the more mesenchymal Y and N cell lines ^47, 48^. Integrin alpha 6, which binds laminin when partnered with beta1 integrin, was detected in all human SCLC lines assessed, except SCLC-N^H446^ **(Figure 5 A, B)**. Laminin gamma 1 was present in all human SCLC lines assessed, except SCLC-A^H889^, and was most highly expressed in the two Y cell lines along with the A cell line H69. Interestingly, similar to mouse non-NE^Hes1-GFP+^ cells, fibronectin is highly abundant in SCLC-Y^H196^ cells; however, it is below the level of detection in the SCLC-Y^H841^ cell lysates **(Figure 5 A, B)**. Overall, these data are consistent with general enrichment of integrins and ECM proteins in the non-NE Y subtype in both human and mouse SCLC.

## Discussion

Secreted factors from non-NE SCLC cells have been shown to support the proliferation and survival of NE SCLC cells, both *in vitro* and *in vivo* ^15^. As NE cells are thought to be the major tumor-promoting component ^14^, understanding mechanisms that support their growth and survival are important. To understand better this intercellular communication, we investigated the role of SEVs in this process. We found that conditioned medium and SEVs derived from non-NE cells not only enhance the growth and survival of NE SCLC cells but also induce a morphological shift from cell-cell to cell-substrate adhesion. Proteomic analysis of the non-NE EVs revealed that multiple ECM and adhesion proteins are enriched cargoes, including laminins associated with stem cell behavior and stromal matrix molecules such as fibronectin and tenascin C. Several, but not all of the ECM proteins, reproduced the growth and adhesion promoting behavior of conditioned media and SEVs. Human non-NE SCLC cells also expressed higher levels of adhesion and ECM molecules than NE SCLC cells. These data suggest that a key role of non-NE SCLC cells and the EVs that they secrete is to provide a supportive niche for NE SCLC cells through providing key ECM growth, adhesion, and survival signals.

SCLC is thought to originate primarily from specialized sensory pulmonary neuroendocrine cells (PNECs) in the lung epithelium ^11^. Loss of RB and p53 are the most frequent genomic alterations in SCLC and targeted inactivation of those genes in PNECs leads to the efficient formation of SCLC tumors in mice ^11^. Ouadah *et al.* reported that PNECs with Rb and p53 deletions grow slowly and have the ability to self-renew and act as stem cells ^49^. They also found that Notch signaling is activated in PNECs in response to injury as they are transitioning to become additional cell types. Similarly, we previously found that Notch activation marks the non-NE subtype of SCLC, suggesting that this subtype may share some features with the transit-amplifying cells found in repopulating PNEC stem cell niches ^15^. This non-NE SCLC phenotype is frequently found intermixed with NE cells in human SCLC tumors^13, 15, 50, 51^, suggesting that the exchange of EVs that we describe between mouse non-NE and NE SCLC cells may be recapitulated in human tumors.

The interplay between stem cells and their niche is important for stem cell maintenance and proliferation, as well as context-induced differentiation. The cancer stem cell niche may resemble a perturbed normal niche and include mesenchymal stem and other stromal cells, ECM proteins, immune cells, cytokines, and growth factors ^52–56^. In SCLC tumors, there is a notable low abundance of stromal cells ^57^, and it is thought that non-NE SCLC cells serve as a substitute to provide various support functions. Notably, various ECM proteins, such as fibronectin, laminin, collagen and tenascin, have been reported to protect SCLC cells against apoptosis, chemotherapy and radiotherapy ^58–62^. In addition, SCLC cells plated on ECM-coated surfaces have been previously found to have a flattened appearance ^63, 64^.

A key finding of our study is that non-NE cells synthesize and secrete a large amount of ECM that is carried primarily by EVs. ECM and adhesion receptors such as integrins are frequent cargoes of SEVs in both normal and cancer cell types and have been previously implicated in driving cancer aggressiveness and metastasis ^27, 65–67^. Interestingly, we found that both epithelial (e.g. laminins) and stromal (e.g. fibronectin) types of ECM are enriched on non-NE EVs and function to promote NE SCLC cell adhesion and survival. Western blot analysis of NE^KP3^ cells revealed that the likely integrins that mediate binding to the laminins and fibronectin produced by non-NE cells are respectively alpha 6 beta 1 and alpha 4 beta 1. By contrast, while non-NE^Hes1-GFP+^ cells expressed the laminin-binding integrin alpha 6 beta 1, their integrin repertoire was distinct, with predominant expression of the fibronectin-binding integrins alpha 5 beta 1 and alpha v beta 1 and the collagen-binding integrin alpha 2 beta 1. It is also possible that additional alpha v integrins are expressed.

To explore how the ECM and adhesion molecules that we identified in mouse non-NE and NE cells relate to human SCLC subtypes, we carried out Western blot analysis of cell lines representing 4 SCLC subtypes: SCLC-A, A2, N, and Y ^7, 32^. The most obvious parallel was for integrin alpha 5 subunit, which was strongly expressed in both human and mouse SCLC-Y cell subtypes but not in other subtypes. There was also enrichment of fibronectin in one of the human SCLC-Y cell types, similar to non-NE^Hes1-GFP+^ cells; however, this was not the case for the other SCLC-Y cell line. While the integrin alpha 4 and 6 subunits were most strongly expressed in the mouse NE^KP3^ cells, they were more widely expressed across the hSCLC cell lines, with integrin alpha 6 expressed in all of the subtypes and alpha 4 most strongly expressed in the SCLC-N and SCLC-Y subtypes. One caveat is that we did not yet assess ECM and adhesion receptors for EVs purified from the human subtypes, which may have some variance from the cell line levels. A future direction is to assess adhesion cargoes in EVs purified from human SCLC subtypes and test their role in mediating survival and adhesion. Another future direction is to assess the role of non-NE SCLC EVs in mediating tumor growth *in vivo.*

In summary, we found that EVs from non-NE mouse SCLC cells carry an abundance of adhesion and ECM cargoes that promote adhesion and survival of NE mouse SCLC cells. These data suggest a key interchange in heterogeneous SCLC tumors that may promote tumor growth and aggressiveness.

## Acknowledgments

We thank the members of the Vanderbilt CSBC U54 Research Group for their helpful input. This study was supported by funding from NIH U54 CA217450 and NIH U01 CA224276.

## Supplemental Figures

**Supplemental Figure 1.**
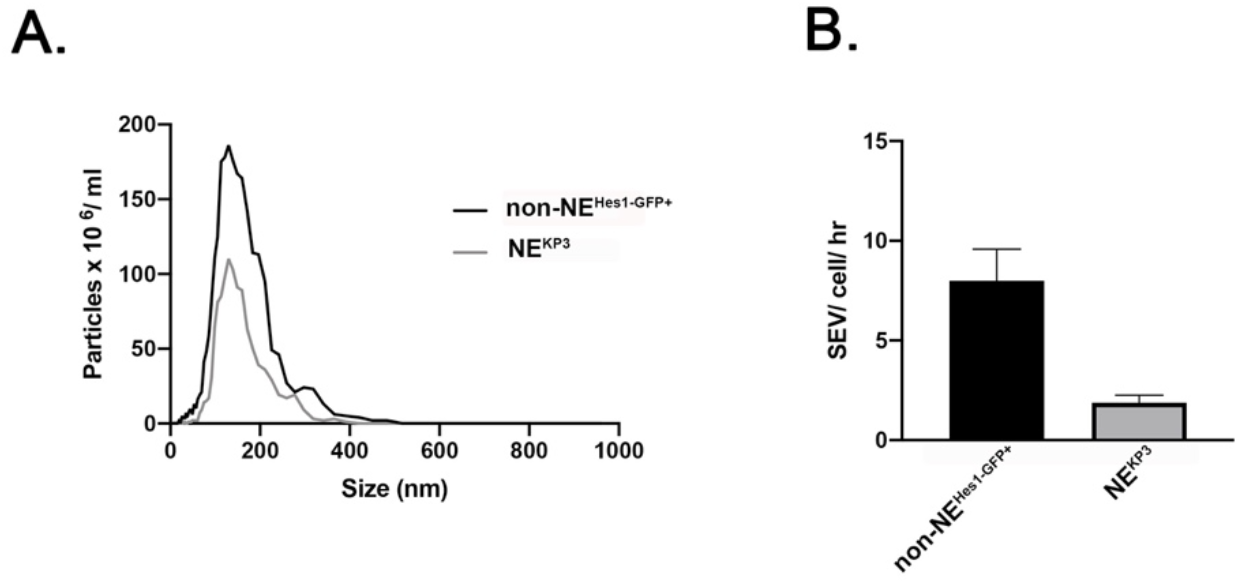
Characterization of mouse SCLC SEVs. A. Representative traces from nanoparticle tracking analysis of SEVs purified for non-NE^Hes1-GFP+^ and NE^KP3^ parental lines. B. Quantitation of SEVs numbers from non-NE^Hes1-GFP+^ and NE^KP3^ parental lines determined in nanoparticle tracking analysis from ≥3 independent experiments.

**Supplemental Figure 2.**
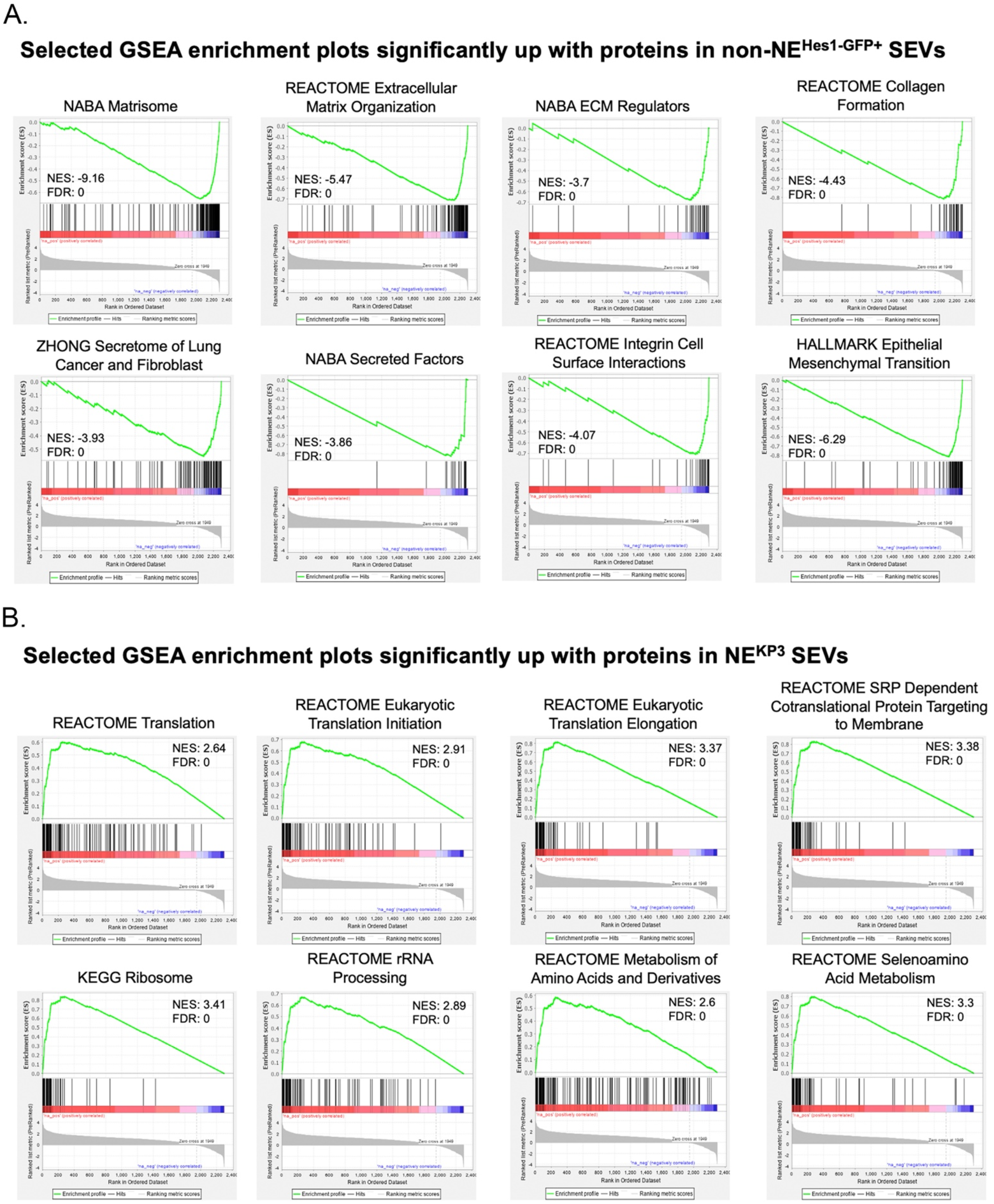
GSEA enrichment plots. A. Selected GSEA enrichment plots significantly up with proteins in non-NE^Hes1-GFP+^ SEVs. B. Selected GSEA enrichment plots significantly up with proteins in NE^KP3^ SEVs. For each plot, the normalized enrichment score (NES) and false discovery rate (FDR) q value are included.

**Supplemental Figure 3.**
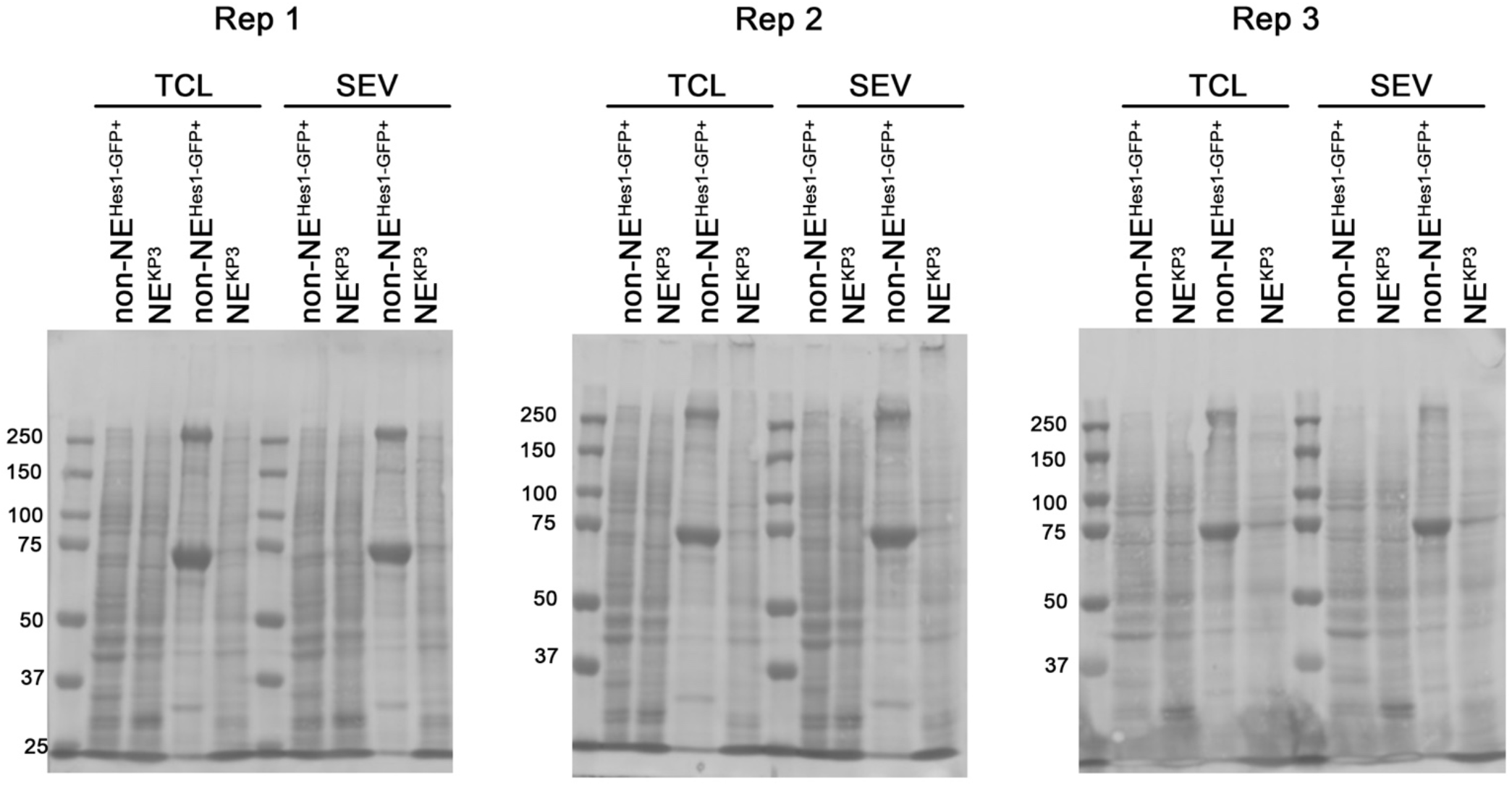
Ponceau-stained membranes from Figure 3. Representative membranes from the Western blot analysis of non-NE^Hes1-GFP+^ and NE^KP3^ total cell lysates (TCL) and small EVs (SEVs).

## Supplemental Tables

**Supplemental Table 1.** Proteins identified in the TMT proteomic analysis

**Supplemental Table 2.** Top 50 GSEA gene sets for non-NE^Hes1-GFP+^ SEVs (first tab) and NE^KP3^ SEVs (second tab). non-NE^Hes1-GFP+^ SEVs are highly enriched in gene sets of proteins associated with epithelial mesenchymal transition (EMT), extracellular matrix organization, matrisome, and integrin cell surface interactions. NE^KP3^ SEVs are enriched in gene sets of proteins associated within ribosome, RNA processing, and translation. For each gene set, the gene ontology (GO) name, # of proteins in that gene set, enrichment score, normalized enrichment score, nominal (NOM) p value, false discovery rate (FDR) q value, and family wise error rate (FWER) p value are listed.

